# Strelka2: Fast and accurate variant calling for clinical sequencing applications

**DOI:** 10.1101/192872

**Authors:** Sangtae Kim, Konrad Scheffler, Aaron L Halpern, Mitchell A Bekritsky, Eunho Noh, Morten Källberg, Xiaoyu Chen, Doruk Beyter, Peter Krusche, Christopher T Saunders

**Affiliations:** Illumina, Inc., 5200 Illumina Way, San Diego, CA 92122, USA.; Illumina Cambridge Ltd., Chesterford Research Park, Little Chesterford, Essex CB10 1XL, UK.; Present address: Seven Bridges Genomics, 101 Euston Road, London NW1 2RA, UK.; Department of Computer Science and Engineering, University of California San Diego, La Jolla, CA 92023, USA.

## Abstract

We describe Strelka2 (<u>https://github.com/Illumina/strelka</u>), an open-source small variant calling method for clinical germline and somatic sequencing applications. Strelka2 introduces a novel mixture-model based estimation of indel error parameters from each sample, an efficient tiered haplotype modeling strategy and a normal sample contamination model to improve liquid tumor analysis. For both germline and somatic calling, Strelka2 substantially outperforms current leading tools on both variant calling accuracy and compute cost.

---

Whole-genome sequencing is rapidly transitioning into a tool for clinical research and diagnosis, a shift which brings new challenges for sequence analysis methods. While there has been considerable progress in developing methods to improve germline and somatic small variant calling accuracy in research applications^1–6^, such methods can be further improved in many respects for the clinical whole-genome sequencing scenario. These improvements include reducing the compute cost/turn-around time of whole-genome analysis, further increasing indel calling accuracy, automating parameter tuning without expert user intervention, and reducing multiple indicators of call quality to a single confidence score for variant prioritization. Here we describe Strelka2, a variant calling method building upon the innovative Strelka somatic variant caller^7^, to improve upon these aspects of variant calling for both germline and somatic analysis. We demonstrate that Strelka2 is both more accurate and substantially faster when compared to current best-in-class small variant calling methods.

Strelka2 germline and somatic analyses share a common series of high-level stages, including parameter estimation from sample data, candidate variant discovery, realignment, variant probability inference, and empirical re-scoring/filtration. The composition of these steps is described in more detail for each type of analysis in Supplemental Fig. 1.

Strelka2’s germline analysis introduces a novel step to adaptively estimate indel error rates from preliminary allele counts in each sample, using a mixture model to estimate both indel variant mutation rates and indel noise rates from a set of error processes (Supplemental Fig. 2). This mixture approach mitigates the impact of context-specific indel error rate variation on variant call accuracy and obviates the need to specify a prior set of common population variants.

Similar to previous work^2,3,5,6^, Strelka2’s germline analysis models haplotypes to provide read-backed variant phasing and reduce the impact of sequencing noise, incorrect read mapping and inconsistent alignment. Strelka2’s haplotype model uses an efficient tiered scheme for haplotype discovery, combining the advantages of a simple model based on input read alignments^3^, and a more complex model using local assembly^2,5,6^, where the appropriate method is selected based on the properties of each variant locus. This tiered haplotype modeling approach is essential to optimize runtime without precision loss. The haplotype model also introduces a novel heuristic filter for sequencer phasing noise, improving the caller’s robustness to a wider variety of potential sequencing artifacts.

In Strelka2’s germline and somatic variant probability models, additional runtime improvements are made in the computation of read likelihoods by enumerating a small number of candidate alignments and using the maximum alignment-specific likelihood to approximate the marginal likelihood, avoiding the computational cost of a complete implementation such as a pair HMM^8^. Within the somatic variant probability model, the original Strelka method has been redesigned with a further novel innovation to account for contamination of tumor cells in the matched normal sample such that somatic recall is improved, especially for liquid tumor analysis. Consistent with the emphasis on automated sample adaption in Strelka2, the liquid tumor model is an expansion of the model’s state space applied to all cases, and thus does not require prior knowledge of the normal sample contamination level.

For both germline and somatic calling workflows, the variant probability model is supplemented by a final empirical variant scoring (EVS) step, motivated in part by machine learning-based variant classification approaches^9,10^. This step uses a random forest model trained on numerous features indicative of call quality to improve precision by accounting for error phenomena that are not adequately represented in the generative variant probability model. Strelka2’s EVS models are pre-trained on data from a variety of sequencing conditions to improve robustness, and produce a single aggregate score which can be used to set application-specific precision levels or prioritize variants for follow-up.

To assess its germline calling performance, we ran Strelka2 on the recent PrecisionFDA Consistency and Truth challenge data^11^ and compared its results with the challenge submissions (Fig. 1a). This comparison shows that for the noisier sequence datasets in the Consistency challenge, Strelka2’s indel accuracy is remarkably higher than the winning challenge submissions, improving upon the indel F-score of the best challenge submission by 3.1%. For the other two Truth challenge data sets with lower sequencing noise, Strelka2 still improves upon the best challenge submission with an indel F-score improvement of 0.08%. For single nucleotide variants, Strelka2 gave competitive results within only 0.05% - 0.1% of the best submissions (Supplemental Fig. 3). These results are striking when considering that all Strelka2 analyses used default parameters, a single read mapper and no input from population variant databases, whereas the top results of the PrecisionFDA challenge were obtained using pipelines specially trained for the challenge data or by combining results from multiple read mappers and variant callers.

**Figure 1.**
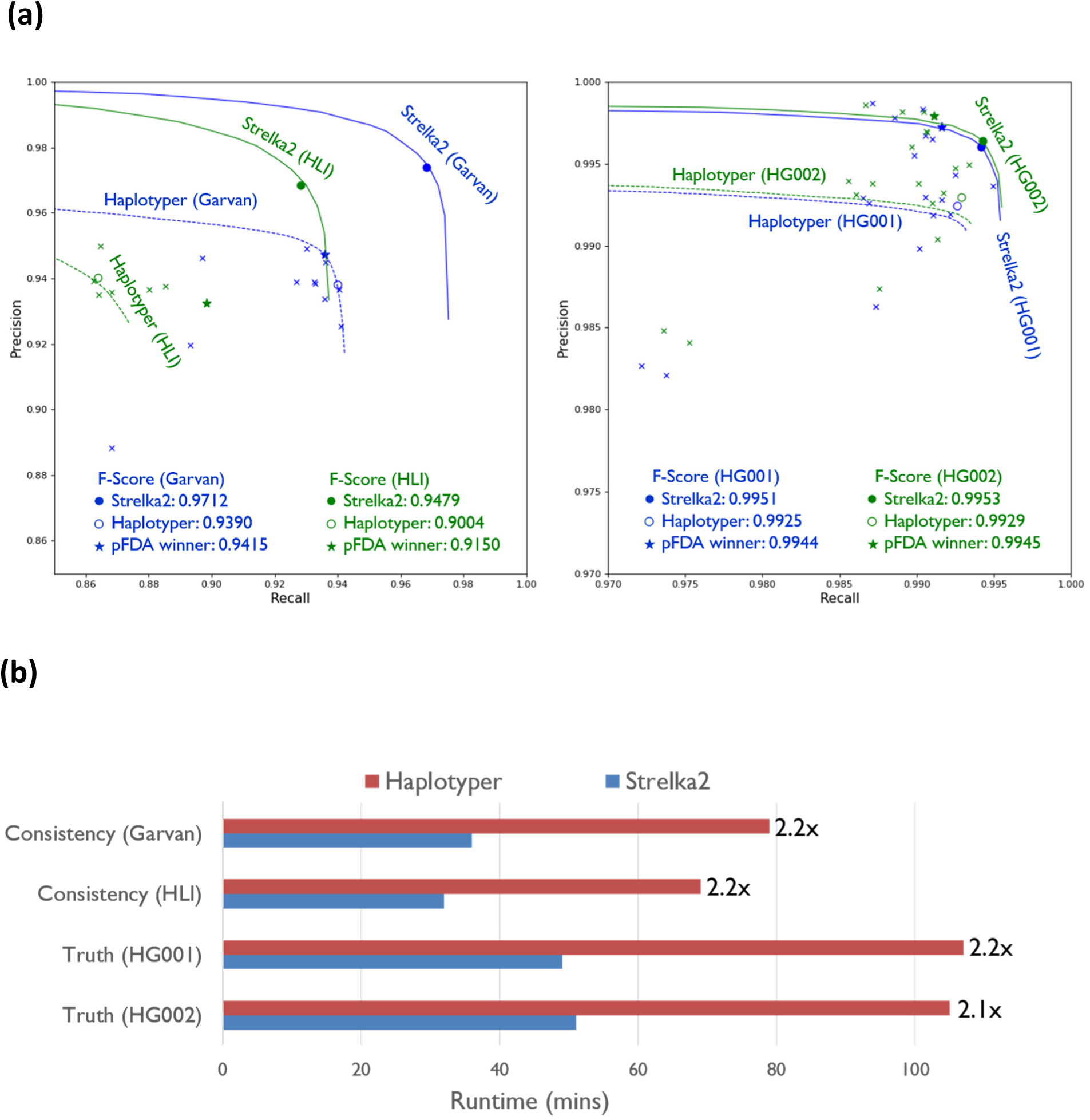
Germline variant calling accuracy and runtime. **(a)** Germline indel calling accuracy of various pipelines, separately plotted for the Consistency (left) and Truth (right) challenge datasets. Filled and empty circles denote passing calls from Strelka2 and Sentieon DNAseq Haplotyper, respectively. Crosses represent passing calls from PrecisionFDA submissions and stars denote the submissions with the best F-scores. For all 4 datasets from these challenges, we mapped each sample using bwa-mem, and ran Strelka2 on default settings. We then compared results against the latest genome in a bottle truth set^15^ using hap.py^16^ (see **Supplementary Note 1** for details). **(b)** Runtime for Strelka2 and Haplotyper for the 4 PrecisionFDA datasets measured on the same compute hardware with two Indel Xeon E5-2680 v4 CPUs (total 28 cores). The coverages of the datasets are 40x, 35x, 50x, and 50x for Consistency (Garvan), Consistency (HLI), Truth (HG001), Truth (HG002), respectively.

To assess runtime, we benchmarked Strelka2 against a recently released high-speed GATK Haplotyper reimplementation (Sentieon DNAseq Haplotyper) that is over 10x faster than the original HaplotypeCaller^12^. On the PrecisionFDA datasets discussed above, Strelka2 was 2.1 times faster than Sentieon DNAseq Haplotyper on average on the same computer hardware while also outperforming it in accuracy, with an average F-score improvement of 2.1% for indels and 0.29% for SNVs (Fig. 1).

We evaluated Strelka2’s somatic variant calling accuracy by mixing sequencing data of unrelated individuals to simulate impure tumor and matched normal samples. For this purpose, we used NA12878 and NA12877 to represent, respectively, the tumor and normal samples. We simulated datasets with tumor purities of 20%, 50%, and 80%, and one matched normal sample with 90% purity. The truth set for these evaluations were the Platinum Genomes^13^ variants in NA12878 where the corresponding NA12877 genotype is homozygous reference. Using the in-silico mixtures, we compared the somatic variant call accuracy of Strelka2 with a recent high-speed MuTect2^4^ reimplementation (Sentieon TNseq TNhaplotyper). As summarized in Fig. 2a, Strelka2 shows substantially higher precision than TNhaplotyper at all recall thresholds over all test datasets, with an average F-score improvement of 29% for SNVs and 35% for indels. We note in particular Strelka2’s superior tolerance to normal sample impurity, reflecting updates in Strelka2’s somatic calling model to better support such contamination in liquid and late-stage solid tumor analyses. This was tested using the 80% purity tumor sample and noting the impact on somatic F-score when the normal sample purity changed from 100% to 90%. Strikingly, for TNhaplotyper the F-score dropped from 77% to 30% for SNVs and from 47% to 17% for indels. For Strelka2, the impact was substantially smaller, changing from 96% to 90% for SNVs and from 82% to 65% for indels.

**Figure 2.**
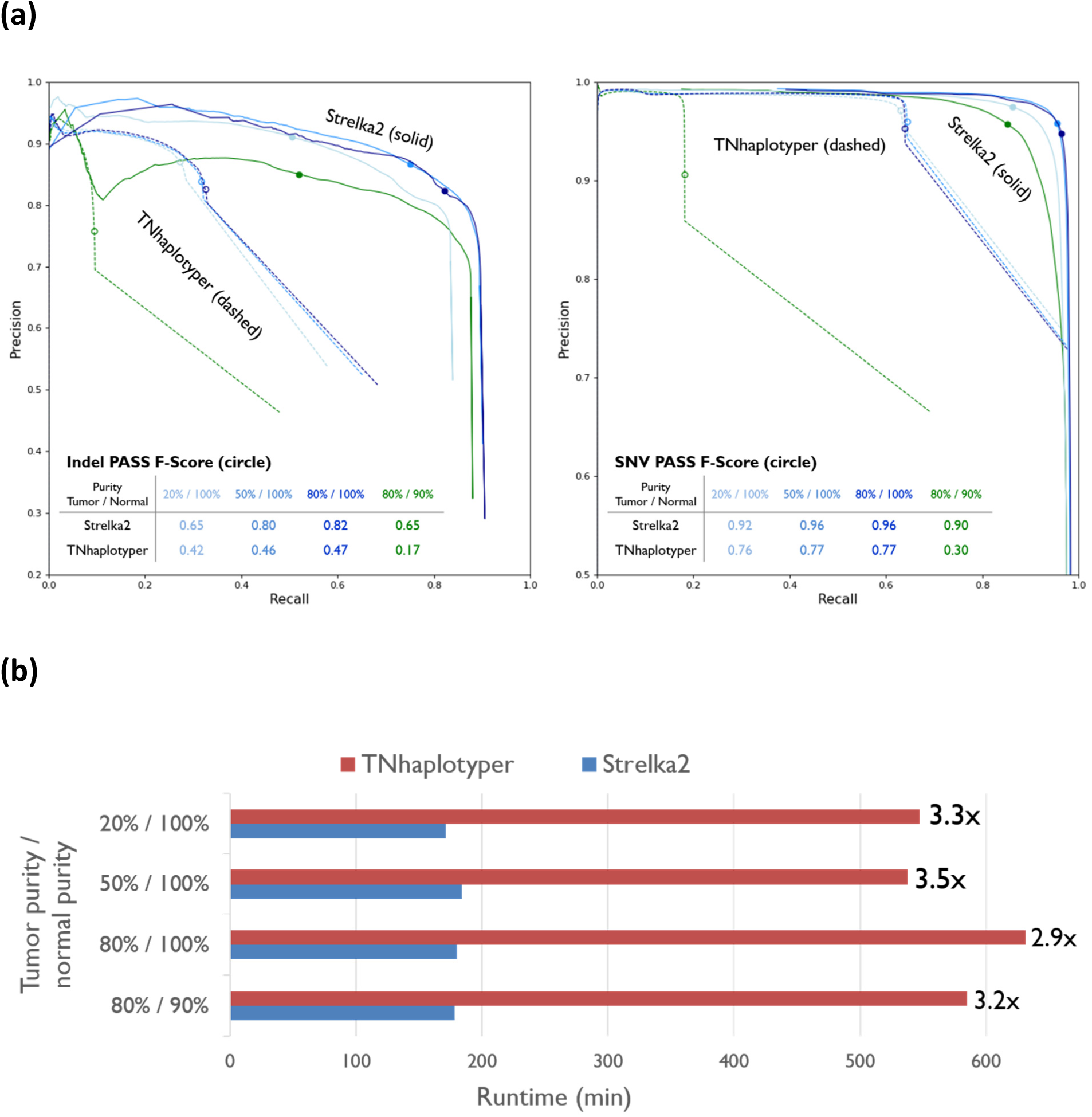
Somatic variant calling accuracy and runtime. **(a)** Somatic variant calling accuracy for Strelka2 and Sentieon TNseq TNhaplotyper, separately plotted for indels (left) and SNVs (right). Datasets are denoted by x / y, where x and y represent tumor (NA12878) and normal (NA12877) purity, respectively. Filled and empty circles denote the passing calls from Strelka2 and TNhaplotyper, respectively. We mapped each sample using bwa-mem, and ran Strelka2 on default settings. Results were compared against the truth set consisting of the variant calls in NA12878 where the corresponding NA12877 genotype is homozygous reference (see **Supplementary Note 1** for details). **(b)** Runtime for Strelka2 and TNhaplotyper for the 4 admixture datasets, measured on the same compute hardware with two Indel Xeon E5-2680 v4 CPUs (total 28 cores). For each dataset, the coverages of tumor and normal samples are 110x and 37x, resepectively.

We assessed runtime for Strelka2’s somatic analysis and found that, as for the germline analysis, Strelka2 is substantially faster than available alternatives. For the above somatic analysis using in-silico sample mixtures, Strelka2 demonstrated an average runtime advantage of 3.2x over TNhaplotyper, which itself is over 10x faster than the original MuTect2 implementation (Fig. 2b)^12^.

In the above analyses, we demonstrate the effectiveness of multiple statistical modeling and algorithmic innovations in Strelka2, resulting in remarkable improvements to accuracy and runtime for both germline and somatic calling. We reiterate that results were generated with default method settings appropriate for factory-scale analysis, not requiring human intervention to parameterize or connect complex sequences of tools. All results use only a reference genome and one alignment file per sample as input. Additionally, no prior variant databases are used in the calling process, reducing the potential for bias against rare variants or ancestry-dependent artifacts.

Improvements to Strelka2 continue in several areas. We have already generalized Strelka2’s germline analysis for RNA-Seq (not described here) and efforts are ongoing to improve mitochondrial variant calling and to integrate with structural variant predictions. Generalization of Strelka2’s adaptive indel error estimation methods to mitigate the impact of context-sensitive base-calling errors on SNV calling has been prototyped and shows promise. The application of these techniques to somatic variant analysis, while considerably more challenging, could substantially improve our ability to call very low-frequency variants. We see such adaptive parameterization improvements as complementing rather than competing with recent trends emphasizing a greater focus on empirical machine learning approaches to variant calling. Indeed, the improvement of generative sequencing error models to more closely represent the sample data should sharpen the effectiveness of downstream machine-learning approaches by reducing confounding error terms, a circumstance we have already leveraged to improve the accuracy of Strelka2.

## ACKNOWLEDGMENTS

We thank Eddy Kim and Ying Wei for testing Strelka2, and Semyon Kruglyak, Ben Moore, Jared O’Connell, and Efstanthios Kanterakis for helpful discussions and comments.

## AUTHOR CONTRIBUTIONS

All authors designed the algorithms and implemented the Strelka2 software. S.K., and C.T.S. designed and performed the analyses. S.K., K.S., C.T.S. wrote the manuscript with input from all authors.

## Online Methods

### Parameter estimation

#### Chromosome depth estimation

An initial step in all workflows is the rapid estimation of the sequencing depth for each chromosome, which for somatic analysis is computed only for the normal sample.

#### Indel error model

Indel sequencing errors are modeled in the variant calling steps below as a process which occurs independently in each read, with some fixed probability of an indel error occurring as a function of the short tandem repeat (STR) context (Supplemental Fig. 2). For germline variant calling, these error probabilities are estimated from the sequencing data of each input sample in two steps. First, mapped sequencing data are analyzed at a subset of sites across the genome to produce error counts for various sequencing contexts. Second, the counts are used to estimate the parameters of interest. For somatic variant calling, a simpler non-adaptive approach is used in which the indel error parameters are pre-set based on empirical observation of indel calling performance and error rates are a function of the homopolymer context length *r* only.

#### Error counting

At every counted site in the genome, the number of reads supporting each potential allele are accumulated by *context*. The counting process uses a read realignment strategy similar to that used by the variant calling process explained below.

Each STR tract with pattern size *s* and repeat count *r* at locus l is counted as a single observation for the context {*s*, *r*} and the resulting locus count vector *c*_*l*_ for that observation (with elements *c*_*l*_ (*y*), *y* ∈ *Y*, corresponding to the observed set of alleles *Y*) is included in the set of locus count observations *C* (*s*, *r*).

#### Error rate estimation

Every indel locus is modeled as belonging to either a *clean* state (generating essentially no indel errors) or a *noisy* state (generating indel errors independently across reads according to a set of error probabilities to be estimated), with the overall error probabilities being drawn from the resulting two-state mixture model. The allele counts, in turn, are modeled as drawn from a mixture over possible genotypes, with the genotype-specific distributions being multinomial for homozygous genotypes and mixtures of two multinomials for heterozygous genotypes. The multinomial distributions are governed by the local coverage *X* = ∑_*Y*_ *c*(*y*) and by rates selected from the vector of available error rates according to the alleles *h*_1_ and *h*_2_ comprising the genotype (Supplemental Fig. 2a).

For every STR context {*s*, *r*} we define the following parameters. Together, the *e* parameters comprise *E*(*s*, *r*):

- *p*_*n*_ (*s*, *r*): probability of being in the noisy state
- *e*_*i*_ (*s*, *r*):noisy-state probability of insertion error resulting in a non-reference variant
- *e*_*d*_ (*s*, *r*): noisy-state probability of deletion error resulting in a non-reference variant
- *e*_ref_ (*s*, *r*): noisy-state probability of indel error resulting in reversion to reference
- *e*_*i*,clean_ (*s*, *r*): clean-state probability of insertion error resulting in a non-reference variant
- *e*_*d*,clean_ (*s*, *r*): clean-state probability of deletion error resulting in a non-reference variant
- *e*_ref,clean_ (*s*, *r*): clean-state probability of indel error resulting in reversion to reference
- ***θ***(*s,r*): probability that a locus with the given context has an indel allele.

We fix all of the clean-state error probabilities to a small constant value: 1×10^-8^ and set *e*_ref_ to 0.01during parameter estimation. During variant calling (described below), the value of *e*_ref_ is a function of the corresponding *e*_i_ or *e*_d_. To improve robustness, the indel mutation rates *θ* were pre-estimated from all autosomes of a fixed human training sample and set as constant values in the workflow. The remaining parameters are estimated on a per-sample basis by maximizing the likelihood of the observed counts:

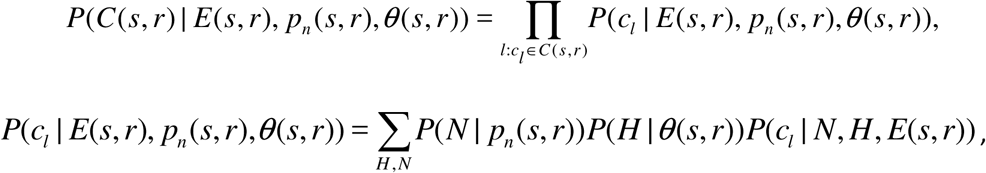

Where 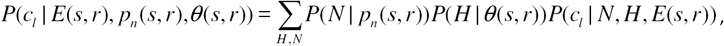

H = (*h*_1_, *h*_2_) is a variable indicating the specific allele hypotheses under consideration and *N* ∈{noisy, clean} is a variable indicating whether the observation was generated by the noisy or the clean state. For each possible genotype *G* ∈{*g*_homref_, *g*_het_, *g*_homalt_, *g*_hetalt_} we consider only one hypothesis H, obtained by finding the two most likely (by number of supporting counts) non-reference indel alleles *y*_1_ and *y*_2_ and setting

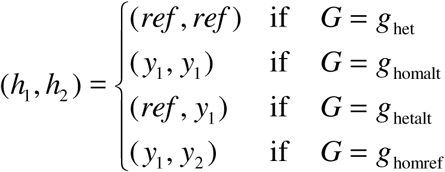

The noisy-state prior is *P*(*N* = *noisy*|*s, r*) = *p_n_*(*s, r*); *P*(*N* = *clean|s, r*) = 1-*p_n_*(*s,r*) and the genotype prior *P*(*G* | θ (*s*, *r*)) is defined in the Germline Probability Model below.

For an allele *y* other than *ref*, *y*_1_, and *y*_2_ assign the error probability *e*(*y*) *e*_*i*_(*s*, *r*), *e*_*d*_ (*s*, *r*), *e*_*i*,clean_(*s*, *r*), or *e*_*d*,clean_ (*s*, *r*) according to the value of *N* and whether *y* is an insertion or deletion with respect to the reference allele. For *y* = *ref*, assign *e*(*y*) = *e*_ref_ (*s*, *r*) or *e*(*y*) *e*_ref,clean_ (*s*, *r*) according to the value of *N*. The probabilities *e*(*y*_1_) and *e*(*y*_2_) of errors resulting in *y*_1_ and *y*_2_ respectively do not need to be assigned specific values due to the approximation below. Finally, let 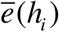 be the probability with which *h*_*i*_ is sequenced correctly, with 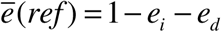 in the noisy state and 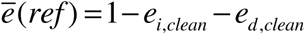 in the clean state.

When *h*_1_ = *h*_2_ (homref or homalt genotype), the likelihood of the count vector is:

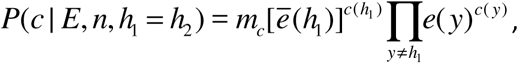

where *m*_*c*_ = (∑_*y*_ *c*(*y*))!/ ∏_*y*_(*c*(*y*)!) is the corresponding multinomial coefficient. Since this coefficient does not depend on any model parameters, it is ignored during parameter estimation.

When *h*_1_ ≠ *h*_2_ (het or hetalt genotype), the likelihood is:

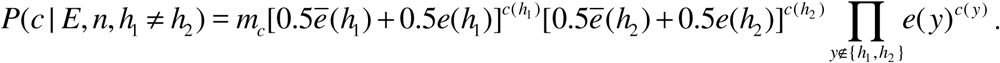

Approximating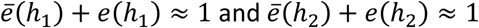, this simplifies to:

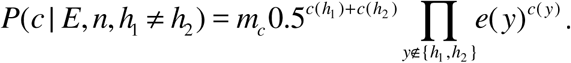

To reduce the number of estimated parameters, we model homopolymer repeats (*s* = 1) with repeat counts 2 *≤ r ≤* 16 and dinucleotide repeats (*s* = 2) with 2 *≤ r ≤ 9* as log-linear in *r*, allowing us to estimate values for (*s, r*) *∈* {(1,1), (1,2), (1,16), (2,2), (2,9)} and interpolate between these values. The values at (1,16) and (2,9) are used for *r* > 16 or *r* > 9 respectively.

When using the estimated parameters for variant calling (Supplemental Fig. 2b, described below), we assume that all sites at which candidate haplotypes have been generated belong to the noisy state, so that the mixture model formulation is not needed. For this reason, only the noisy-state error probabilities are passed on for downstream use. We also fix the insertion and deletion error rates to be used for calling to the geometric mean of the insertion and deletion estimates for each STR context.

### Candidate variant discovery

#### Input read processing

The input alignment files are scanned for reads. Reads are filtered out if they are marked as not passing primary analysis filters, PCR/optical duplicates, unmapped, secondary or supplemental. Indels in the remaining reads are left-shifted and normalized.

#### Tiered haplotype model

Each sample’s ploidy introduces constraints that can be used to reduce errors due to sequencing noise, incorrect read mapping, and inconsistent alignment. In a simple haplotype model, such as that described in FreeBayes^3^, candidate haplotypes can be identified from existing read alignments. More advanced haplotype generation methods are less dependent on the input read alignments, often using local assembly to identify longer consensus haplotypes, such as the haplotype generation methods used in Platypus^5^, Scalpel^6^ and GATK HaplotypeCaller^2^, among others. Strelka2’s germline caller uses a *tiered haplotype model* where a fast alignment-based approach is used to handle simpler variant loci, and an assembly-based approach is selected to improve accuracy in more complex cases.

The haplotyping steps are: detecting short clusters of sequence variation called *active regions*, generating candidate haplotypes in active regions, filtering candidate haplotypes to reduce noise, and discovering primitive SNVs and indels. Haplotyping is currently performed independently for each sample and is not available for somatic variant calling.

#### Active region detection

To detect active regions, we first identify loci that are likely to have variants, which we call *variant loci*. To identify variant loci, for each locus we calculate a variant evidence score while reading alignments as follows: a mismatch at locus *i* increases the score at *i* by 1, an insertion between locus *i* and *i* +1 increases the scores at *i* and *i* +1 by 4, a deletion of loci [*i*, *j*] increases the scores in [*i* -1, *j*] by 4, and a soft-clipped segment ending (starting) at locus *i* increases the scores at *i* and *i*+1 (*i* -1) by 4. A locus with a variant evidence score *c* and a coverage *d* becomes a variant locus if (1) *c ≥* 0.35 × *d* or (2) *c ≥* 9 and *c ≥* 0.2 × *d*. Afterwards, nearby variant loci are clustered if they are no more than 13 bases of each other. For the clusters including two or more variant loci, the cluster region is further extended to the surrounding loci so that the first and last locus are not within a homopolymer or STR region. This extension is needed because alignments that do not fully span such repeats are often erroneous and relying on them may lead to generating incorrect haplotypes. To accomplish this, we detect *anchor loci* that are not variant loci and also do not belong to a homopolymer (of lengths no less than 3) or STR (of repeat unit lengths between 2 and 50). Given a cluster of variant loci, the active region is created between the closest anchor loci before and after the first and last variant loci.

#### Haplotype generation

Given an active region of size of 250 or smaller, haplotype generation is attempted using either the alignment-based or assembly-based model. The decision is made based on the fraction of reads that fully cover the active region (called covering reads): the assembly-based model is used if fewer than 65% of all reads that overlap with an active region are covering reads.

If the alignment-based model is selected, then for each covering read, the segment aligned to the active region is extracted as a candidate haplotype. If a candidate haplotype *s* is extracted from a read *r*, we call *r* a supporting read of *s*, such that for each candidate haplotype *s* a set of supporting reads is identified.

If the assembly-based model is selected, local de novo assembly is run using a de Bruijn graph approach similar to that described in TIGRA^14^. Prior to assembly, the target active region [*i*, *j*] is expanded to [*I'*, *j*'] to improve the identification of contigs which span the full locus. *I’* and *j* ‘are chosen as the mininum and maximum value satisfying the following conditions: *i* –9 ≤ *i*’≤ *i, j*≤ j’≤ *j+9* and there is no variant locus in *i’* ≤ *pos* ≤ *i* –1 and *j* +1 ≤ *pos* ≤ *j*’. This expansion allows the indentification assembled contigs which span the full locus by identifying those that share the same prefix (reference segment at [*i’*, *i*]; denoted by a prefix anchor) and suffix (reference segment at [*j*, *j*’]; denoted by a prefix anchor). All the reads that (fully or partially) overlap with the expanded active region are used as input to the assembly procedure. After assembly is finished, only the contigs including both prefix and suffix anchors are selected and the prefix and suffix anchors are removed. Each such contig becomes a candidate haplotype and the set of reads supporting the contig is identified.

Haplotype generation for an active region is considered unsuccessful if the assembly procedure is selected and assembly is unable to generate at least one non-reference candidate haplotype. If haplotype generation does not succeed, indel candidates can still be generated as detailed below without the benefit of haplotyping.

#### Haplotype filtration

If haplotype generation is successful (using either alignment-based or assembly-based methods), candidate haplotypes are ranked by decreasing read support; those with fewer than 3 supporting reads or ranking below the top *x*, for *x* the expected sample ploidy (assumed to be diploid in the current procedure), are excluded from further processing. If there is more than one remaining haplotype, an additional filtration step is applied to reduce candidates produced by phasing noise in the sequencing process across a homopolymer. The test assumes that the candidate haplotype with the highest read support, *h*_1_, is true, and identifies whether the candidate haplotype with next highest level of read support, *h*_2_, is a phasing noise artifact introduced while reading *h*_1_. The conditions which trigger this filter are (1) *h*_1_ and *h*_2_ are the same length with only one mismatching basecall, (2) all reads supporting *h*_2_ are observed on only one strand, and (3) the basecall mismatch between *h*_1_ and *h*_2_ occurs at one of the ends of the sequence, and causes *h*_2_ to contain an uninterrupted homopolymer at least 11 bases long. If these conditions are met, all haplotype candidates besides *h*_1_ consideration.

#### Primitive allele discovery

After filtration, the remaining candidate haplotypes are aligned to the reference and primitive alleles (SNVs and indels of size 50 and smaller) are annotated as *discovered*. These discovered primitive alleles are used to improve SNV and indel calling in downstream procedures.

#### Indel candidacy

Strelka2 uses indel candidacy as a preliminary filter to eliminate indel observations likely to have been generated by error processes. Candidate indels are considered during read realignment and indel genotyping in all samples. To become a candidate, an indel variant must minimally have 2 reads supporting it in at least one sample. If haplotype modeling is enabled, a candidate indel belonging to an active region where haplotyping was successful must also have been discovered through haplotype alignment in at least one sample. If an indel observation satisfies these conditions, Strelka2 evaluates its candidacy status using a one-sided binomial exact test, with the null hypothesis being that the indel coverage is generated by indel error processes. The indel is considered a candidate variant if the resulting p-value is below 1X10-^9^.

### Read realignment and variant probability inference

#### Read realignment

Following the discovery of candidate alleles, reads are realigned to these candidates. This realignment step has two primary functions. The first is to generate the set of most likely alignments under the assumption that the read was generated by a particular candidate haplotype. Such alignments are used to assess the read’s relative support for different indel alleles. The second function is to create a single *representative* alignment to use for SNV calling.

The alignment search uses a starting alignment provided by the input alignment file, as well as a set of intersecting candidate indels. If the read intersects at least one candidate indel, a set of alignments is built from the starting alignment by recursively toggling indels from the candidate set. Each toggling operation produces three alignments: the input alignment itself, and two alignments constructed by adding or removing the indel in question such that the input alignment is unchanged (1) to the left or (2) to the right of the indel. For efficiency, the search recursion is limited to depth 5.

#### Germline Probability Model

At every locus where candidate variant alleles have been proposed, Strelka2 calculates posterior probabilities for a range of hypotheses in each sample. Each germline hypothesis comprises a specific pair of alleles that determine its genotype *G ∈* {*g*homref, *g*_het_, *g*_homalt_, *g*_hetalt_} with the potential genotypes respectively corresponding to non-variants and to variants which are heterozygous, homozygous, and heterozygous with two non-reference alleles.

The posterior probability of a hypothesis *H* conditioned on the observed data *D* and *Q* is:

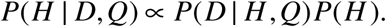

The hypothesis-specific likelihood *P*(*D* | *H*, *Q*) is described under “Shared Probability Model” below. The hypothesis prior depends on the corresponding genotype prior *P*(*G*), defined for variant calls in terms of the prior probability that a chromosome locus is non-reference, *θ* as follows:

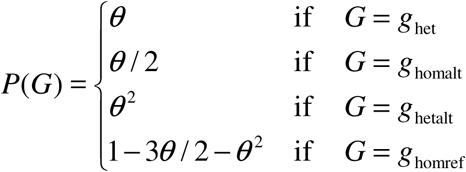

For SNVs, all three possible non-reference alleles are considered at any potential variant site, thus each variant genotype maps to three specific SNV hypotheses with uniform probability: *P*(*H*) = *P*(*G*) 3 when *G ≠ g*_homref_. For indels, up to two non-reference alleles are considered per sample and *P*(*G*) is again divided between matching hypotheses.

#### Germline Variant Phasing

As previously noted, Strelka2 defines an active region around dense variants and infers 2 haplotypes for the region. These haplotypes are used to phase SNVs and indels within the same active region. The phasing is conducted after scoring and genotyping. For each heterozygous variant belonging to an active region, Strelka2 matches the variant alleles to the active region haplotype to appropriately phase the genotype allele order.

#### Somatic Model

The somatic calling model assumes that the samples are diploid. For both SNVs and indels, the normal genotype states are *G ∈* {*g*_ref_, *g*_het_, *g*_hom_}, referring to a non-variant and a variant which is heterozygous or homozygous in the normal sample, assuming no more than one variant allele in this sample. The tumor genotype states are *G ∈* {*g*_nonsom_, *g*_som_}, referring to the absence and presence of a somatic variant in the tumor sample, respectively. The method approximates a posterior probability on the joint tumor and normal genotypes:

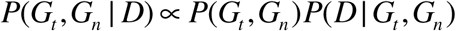

Here *D* refers to the sequencing data from both samples. The likelihood term above is computed by integrating over sample-specific allele frequencies

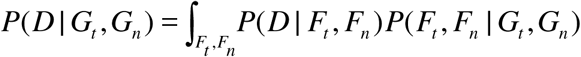

where *F*_*t*_and *F*_*n*_ refer to tumor and normal allele frequencies. The allele frequency likelihood *P*(*D* | *F*_*t*_, *F*_*n*_) is decomposed by sample to *P*(*D*_*t*_ | *F*_*t*_)*P*(*D*_*n*_ | *F*_*n*_), where *D*_*t*_ and *D*_*n*_ indicate tumor and normal sample data. The sample-specific allele frequency likelihoods *P*(*D*_*t*_ | *F*_*t*_) and *P*(*D*_*n*_ | *F*_*n*_) are as described in the Shared Probability Model section below. The genotype prior probability *P*(*G*_*t*_, *G*_*n*_) and the joint allele-frequency distribution *P*(*F*_*t*_, *F*_*n*_ | *G*_*t*_, *G*_*n*_) are detailed in the present section.

The posterior probability over tumor and normal genotypes *P*(*G_t_*, *G_n_* | *D*) is used to compute the *somatic variant probability*.

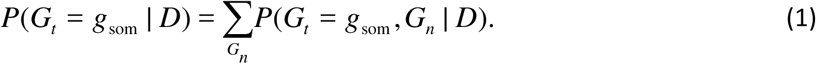

Somatic variant calls are reported jointly with associated calls for the normal sample. For this, we use the joint probability of somatic variation and the maximum likelihood normal sample genotype:

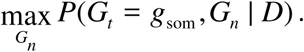

Given the expected rate of variants between two unrelated haplotypes *θ*, the normal sample genotype prior *P*(*G*_*n*_) is

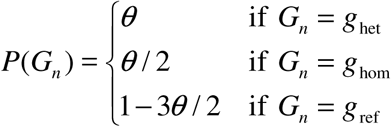

where *θ* is 10^-3^ for SNVs and 10^-4^ for indels. Given the somatic state prior *P*(*G_t_* = som) = *γ*, the joint sample prior is

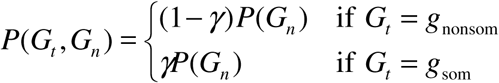

where *γ* is set to 10^-4^ for SNVs and 10^-6^ for indels. These values were chosen empirically to provide reasonable variant probabilities and are not adjusted for different samples in practice.

The prior probability on the tumor and normal allele-frequencies *P*(*F*_*t*_, *F*_*n*_ | *G*_*t*_, *G*_*n*_) encodes the concept that the normal sample is a mixture of diploid germline variation and noise while the tumor sample is a mixture of the normal sample and somatic variation. Let *C*(*F*_*n*_, *G*_*n*_) = 1 if (*F*_*n*_, *G*_*n*_) is (0, *g*_ref_), (0.5, *g*_het_), or (1, *g*_hom_) and *C*(*F_n_*, *G_n_*) = 0 otherwise. The joint frequency prior is then defined as

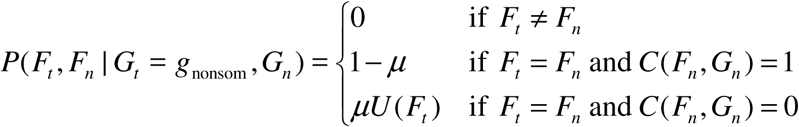

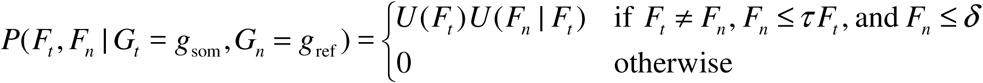

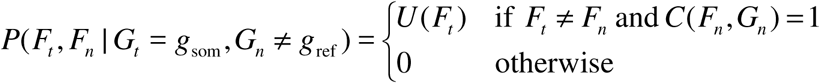

Here, τ and δ represent contamination tolerance terms, *U*(*F_t_*) to a uniform distribution over the allowed tumor allele frequencies, *U*(*F_n_*|*F_t_*) refers to a uniform distribution over the normal allele frequencies satisfying *F_n_* ≤ *τF_t_* and *F_n_* ≤ *δ*, and *μ* indicates the noise term. The contamination tolerance terms are introduced to allow for contamination in the normal sample by some fraction of tumor cells. This is particularly useful for analyses of liquid tumors, where the normal sample may be contaminated by tumor cells. By default, τ and δ are set to 0.15 and 0.05. The noise term abstracts various sequencing, read mapping and assembly issues which could produce an unexpected allele frequency shared in the tumor and normal samples. For SNVs, the noise contribution is set to a constant value *μ*_SNV_ = 5×10^–10^ and for indels it is set as a function of the indel error rate 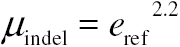

The continuous allele frequencies modeled above are efficiently computed by dividing each allele-pair axis into a set of equidistant points and performing the somatic probability computation over the resulting discrete point set. A resolution of 21 points per axis (i.e., points separated by 0.05) is used for all computations by default.

#### Shared Probability Model

All germline and somatic hypotheses can be generalized as a list of haplotypes *h_i_* with corresponding expected frequencies *f_i_* in each sample. We are interested in the hypothesis specific likelihood for a given sample *P*(*D* | *H*,*Q*) of the sample’s observed set *D* of individual reads *d_j_* (assumed independent) given the observed set *Q* of individual basecall quality scores *q_j_*:

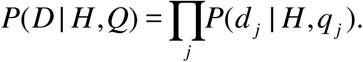

The likelihood for an individual read can be expressed in terms of likelihoods conditioned on each of the potential generating haplotypes:

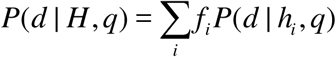

The per-read likelihood *P*(*d* | *h*, *q*) is the probability of an individual read *d*, given its associated basecall qualities *q* and a generating haplotype *h*. In a complete probabilistic implementation (e.g. using a pair HMM), this likelihood would be computed by summing over all possible pairwise alignments *A* in which *d* is aligned to *h*. Strelka2 saves computation by enumerating a small number of candidate alignments and using the maximum alignment-specific likelihood to approximate the marginal likelihood:

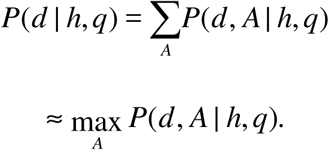

The alignment-specific likelihood scores can be factorized as follows:

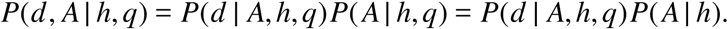

Ignoring possible context effects and accepting the basecall quality scores at face value, the first term is a product of emission scores, *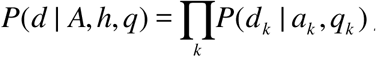*, where emission scores are:

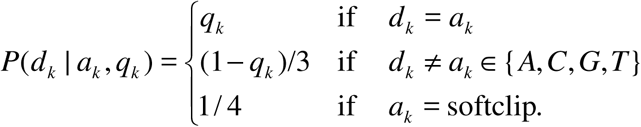

Here, *d*_*k*_ and *q*_*k*_ are the *k* th base in *d* and the corresponding probability of the call being correct (obtained from the basecall quality score) and *a_k_* is the base in *h* to which *d_k_* has been aligned or, if *d_k_* is aligned to an insertion relative to h, the corresponding base of the consensus insertion sequence. The second term is a product of state transition probabilities, using the indel error probabilities *e*_*i*_ (*s*, *r*), *e*_*d*_ (*s*, *r*), and *e*_ref_ (*s*, *r*) described earlier to penalize alignments whenever the read contains an indel with respect to the generating haplotype:

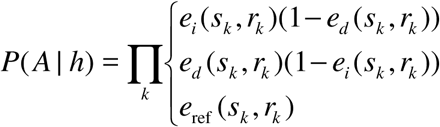

where the product is taken over all positions *k* at which a gap is opened and *reversion to reference* refers to indels that result in the reference allele being generated even though the generating haplotype contained a non-reference allele at position *k*. To compensate for reference-bias in the alignment process, the value of *e*_ref_ is set to a constant factor (1.8) times the corresponding *e*_i_ or *e*_d_. The probabilities calculated in this equation are unnormalized, due to omission of corresponding terms when a gap fails to open; this is corrected by normalizing explicitly during posterior probability calculation.

### Empirical scoring and filtration

The variant calling models (both germline and somatic) provide sufficiently accurate representations of the biology and sequencing process to produce an initial variant probability inference. However, there is additional information not used by the models which is nonetheless predictive of call accuracy. As a final step in the variant calling process, such additional information is extracted as a set of predictive features and used in combination with the probability calculated by the variant calling model to improve call precision. This is done by the Empirical Variant Scoring (EVS) model, a supervised random forest classifier trained on labeled data from sequencing runs performed under a variety of conditions (different sequencers, sample preparation, and coverage). The EVS model provides an aggregate quality score for each variant and allows for convenient exploration of the precision-recall curve.

For each of germline and somatic variant calling, there are two separate feature sets and trained random forest models: one for SNVs and one for indels. In contrast to dynamic re-scoring systems such as the GATK VQSR procedure^9^, the EVS models are pre-trained, allowing Strelka2 to avoid the runtime cost, instability and population variant data requirement of a dynamic approach. When the EVS model is not used, simple cutoffs are applied to a set of features (not necessarily the same set used by EVS) to less precisely filter out potentially problematic calls. For details on EVS training and the full lists of EVS and hard-filter features, refer to **Supplementary Note 2**.

## Data availability

The sequencing data and truth sets used in the germline calling benchmarking are publicly available at <u>https://precision.fda.gov/</u> and <u>https://github.com/genome-in-a-bottle</u>. The sequencing data and truth sets used in the somatic calling benchmarking are available at <u>http://strelka-public.s3.amazonaws.com</u>.

---

**Supplementary Figure 1.**
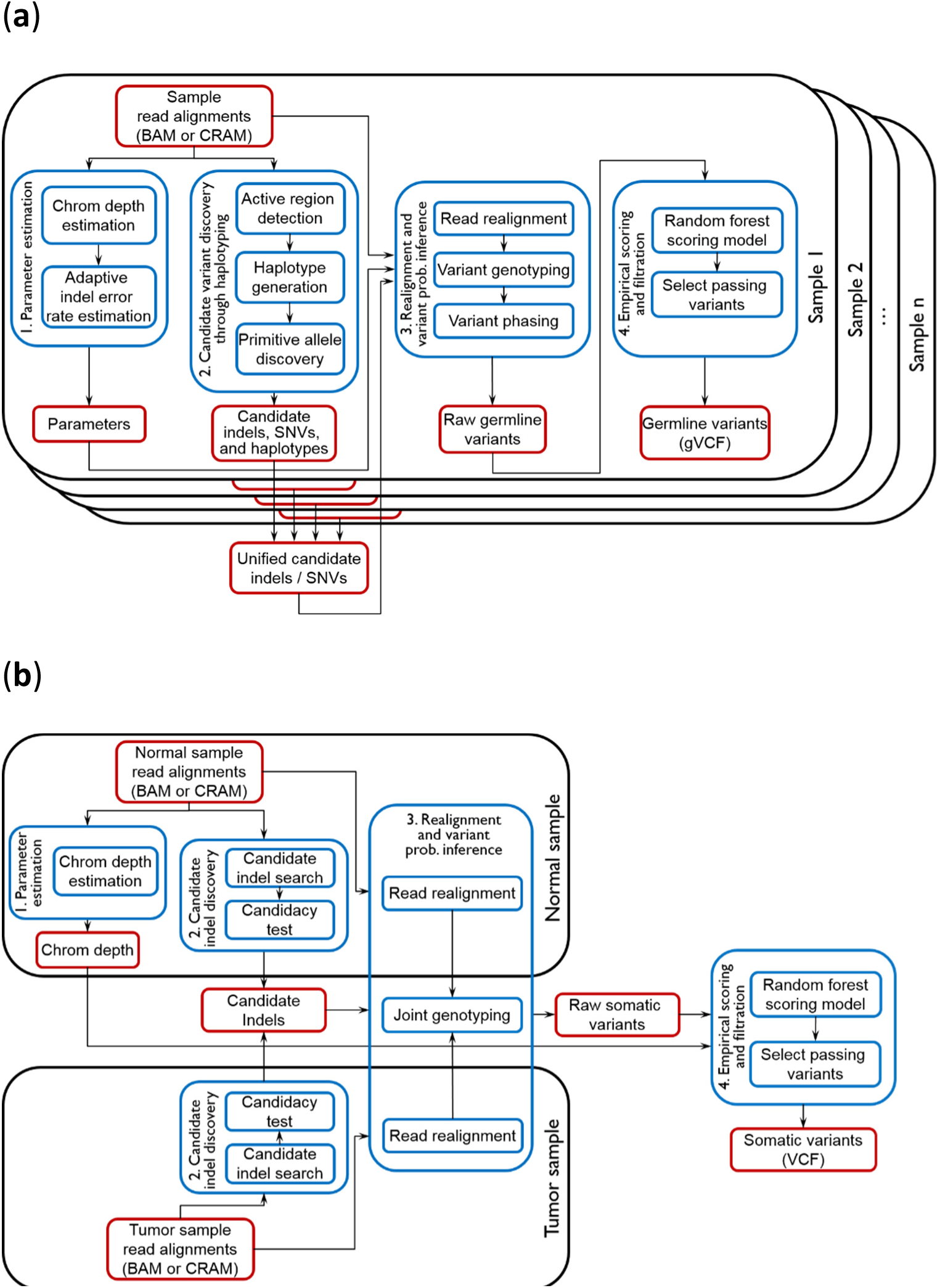
Strelka2 workflows. Strelka2 supports detection of germline variants in small sample cohorts (~10 individuals), and somatic variants from matched tumor-normal sample pairs. These two analyses share several high-level steps, including: (1) parameter estimation, (2) candidate variant discovery, (3) realignment and variant probability inference, and (4) empirical scoring and filtration. Here we diagram an overview of the major workflow components for both **(a)** germline and **(b)** somatic analyses.

**Supplementary Figure 2.**
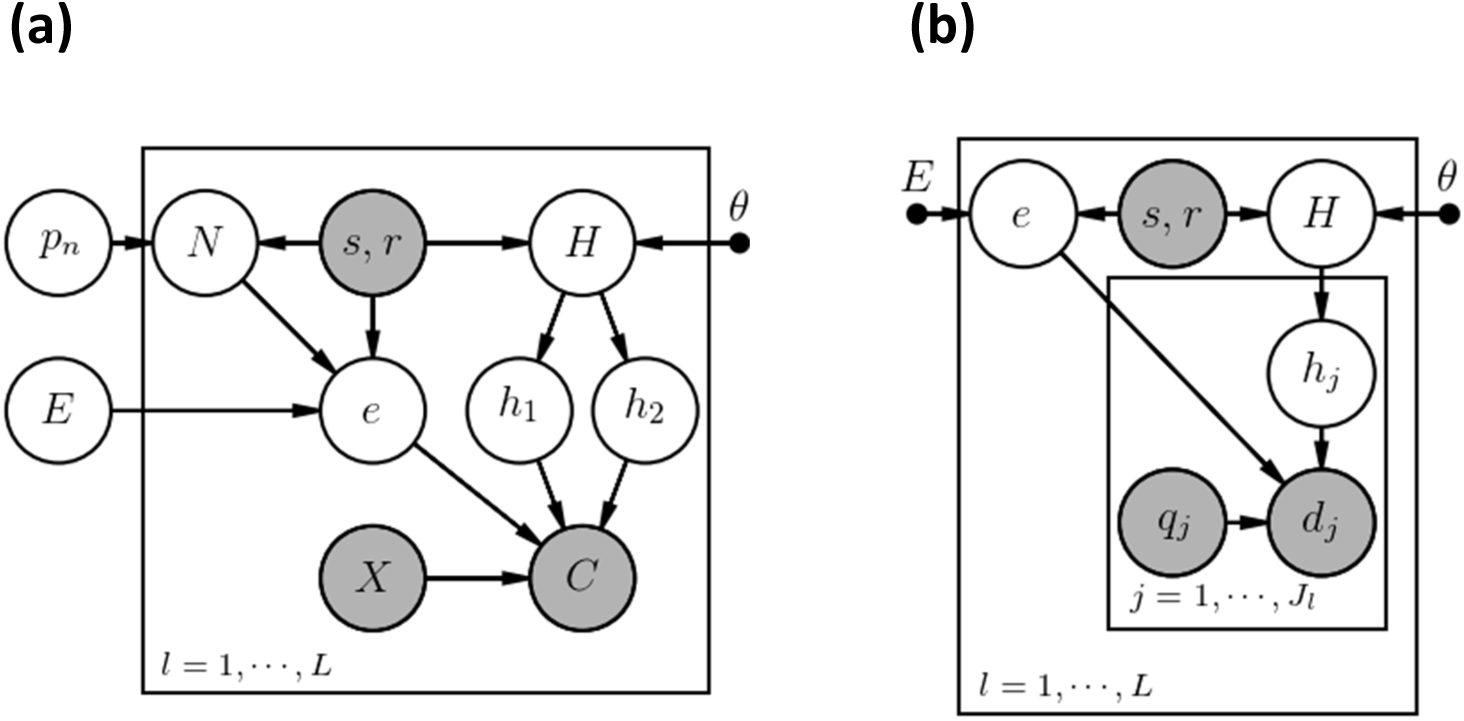
Structure of the germline indel error and variant calling models in probabilistic graphical model plate notation (some details omitted). **(a) Indel error model.** At each locus *l*, a preliminary estimate of the indel allele count vector *C* is modeled as a mixture binomial distribution governed by the two true haplotypes *h*_1_ and *h*_2_ (a function of the unobserved genotype hypothesis *H*), a set of indel error rates *e* (unobserved) and the total count *X* (observed). The error rates are selected from the full set of error parameters *E* according to the sequence context (summarized as an integer pair denoting the size *s* and number *r* of STR repeats; observed) and a binary state variable *N* (unobserved) categorizing the locus as clean (essentially zero error rates) or noisy (prone to indel errors). The genotype *H* and the noisy-clean state variable *N* are drawn from prior distributions that depend, respectively, on a context-specific mutation rate θ shared across samples and a context-specific noisy-state probability *p_n_*. **(b) Variant calling model.** The reads *d_j_* at every locus are modeled as depending on the corresponding base call quality strings *q_j_*, the unobserved haplotype *h*_j_ that generated the read, and the locus-specific error rates *e*. The read-specific haplotype is drawn from the set of haplotypes in the locus-specific hypothesis *H*, of which the prior again depends on a parameter from *θ* according to context. The error rates are again selected from the global vector *E* of error parameters (now treated as fixed), with the difference that all loci analyzed by this model are assumed to be in the noisy state.

**Supplementary Figure 3.**
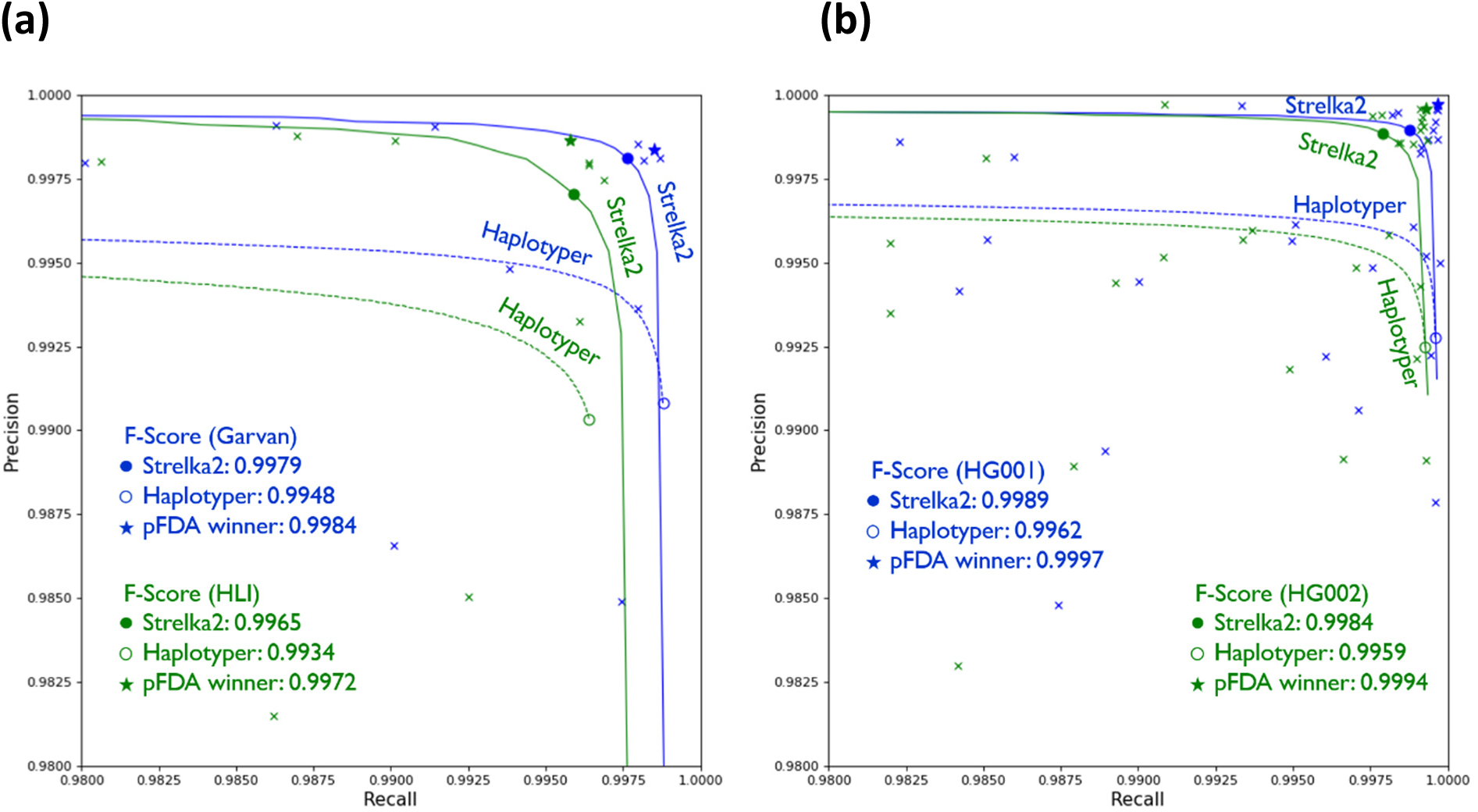
Germline SNV calling accuracy of various pipelines for the Consistency **(a)** and Truth **(b)** challenge datasets. Filled and empty circles denote the precision and recall of passing calls from Strelka2 and all calls from Sentieon DNAseq Haplotyper, respectively. For Haplotyper, the F-scores of passing calls were far lower than those of all calls, so we chose to plot the F-score for all calls. Crosses represent passing calls from PrecisionFDA submissions and stars denote the submissions with the best F-scores.

